# Screening of FDA-approved Drugs and Identification of Novel Lassa Virus Entry Inhibitors

**DOI:** 10.1101/298463

**Authors:** Peilin Wang, Yang Liu, Guangshun Zhang, Shaobo Wang, Jiao Guo, Junyuan Cao, Xiaoying Jia, Leike Zhang, Gengfu Xiao, Wei Wang

## Abstract

Lassa virus (LASV) belongs to the *Mammarenavirus* genus (family *Arenaviridae*) and causes severe hemorrhagic fever in humans. At present, there are no Food and Drug Administration (FDA)-approved drugs or vaccines specific for LASV. Herein, high-throughput screening of an FDA-approved drug library was performed against LASV entry using a pseudo-type virus enveloping LASV glycoproteins. Two hit drugs, lacidipine and phenothrin, were identified as LASV entry inhibitors in the micromolar range. A mechanistic study revealed that both drugs inhibited LASV entry by blocking low-pH-induced membrane fusion. Moreover, lacidipine irreversibly bound to the LASV glycoprotein complex (GPC), resulting in virucidal activity. Adaptive mutant analyses demonstrated that replacement of T40, located in the ectodomain of the stable-signal peptide (SSP), with lysine (K) conferred LASV resistance to lacidipine without apparent loss of the viral growth profile. Furthermore, lacidipine showed antiviral activity and specificity against both LASV and the Guanarito virus (GTOV), which is also a category A new world arenavirus. Drug-resistant variants indicate that the V36M in ectodomain of SSP mutant and V436A in the transmembrane domain of GP2 mutant conferred GTOV resistance to lacidipine, suggesting that lacidipine might act via a novel mechanism other than calcium inhibition. This study shows that both lacidipine and phenothrin are candidates for LASV therapy, and the membrane-proximal external region of the GPC might provide an entry-targeted platform for inhibitors.

## IMPORTANCE

Currently, there is no approved therapy to treat Lassa fever; therefore, repurposing of approved drugs will accelerate the development of a therapeutic stratagem. In this study, we screened an FDA-approved library of drugs and identified two drugs, lacidipine and phenothrin, which inhibit Lassa virus entry by blocking low-pH-induced membrane fusion. Additionally, both drugs extended their inhibition against the entry of Guanarito virus, and the viral targets of lacidipine were identified.

Lassa virus (LASV) is an enveloped, negative-sense, bi-segmented RNA virus belonging to the *Mammarenavirus* genus (family *Arenaviridae*) (1). Mammarenaviruses consist of 35 unique species currently recognized by the International Committee on Taxonomy of Viruses. The original classification of mammarenaviruses, based mainly on virus antigenic properties; serological, genetic, and geographical relationships; and the rodent host, divided them into new world (NW) and old world (OW) mammarenaviruses (2). The OW Lassa virus and some NW mammarenaviruses, including the Junín virus (JUNV), Machupo virus (MACV), Guanarito virus (GTOV), and Sabiá virus (SABV), are known to cause severe hemorrhagic fever and are listed as biosafety level (BSL) 4 agents (3,4). LASV infections cause about 300,000 cases of Lassa fever per year, and the mortality rate in hospitals is reported to be as high as 65–70% (5). At the beginning of this year, a Lassa fever outbreak was reported in Nigeria. From January 1 to March 18, 2018, 376 confirmed cases and 95 deaths have been reported (6).

The LASV RNA genome encodes the viral polymerase, nucleoprotein, matrix protein (Z), and glycoprotein complex (GPC). The GPC is synthesized as an inactive polypeptide and cleaved twice by the signal peptidase and cellular protease subtilisin kexin isozyme-1/site-1 protease, yielding the retained stable-signal peptide (SSP), the receptor-binding subunit GP1, and the membrane fusion subunit GP2 (7-10). The highly conserved arenavirus SSPs contain 58 amino acids that span the membrane twice, with 8 amino acids in the ectodomain, playing essential roles in GPC maturation and downstream functions (11-17). LASV utilizes α-dystroglycan (α-DG) as a primary receptor, and successful infections require the receptor switch to lysosome-associated membrane protein 1 (18-20).

To date, no vaccines or specific antiviral agents against LASV are available. Therapy strategies are limited to the administration of ribavirin in the early course of the illness (21). To address this issue, we screened an FDA-approved drug library of 1018 compounds. The approved drugs have been intensively investigated for safety, pharmacokinetics, and targets; therefore, screening approved drugs for repurposing will increase the speed of discovery and development for treatment (22,23). Drugs targeting viral entry can block replication and spread at an early stage. Since studies of LASV require BSL-4 equipment, we utilized a LASV GPC pseudo-type vesicular stomatitis virus (VSV) containing a *Renill*a luciferase (Rluc) reporter gene for high-throughput screening (HTS) of LASV entry inhibitors, which can be performed in a BSL-2 facility. After three rounds of screening, lacidipine and phenothrin were identified to be highly effective against LASV entry. The hit drugs identified in this study offer potential new therapies to treat arenavirus infections and disease.

## RESULTS

### Screening of an FDA-Approved Drug Library for Inhibitors of LASV Entry

To perform high-throughput screening (HTS) under BSL-2 conditions, we generated a pseudo-type virus bearing the LASV GPC (designated LASVpv) for HTS of entry inhibitors (24). The number of genomic RNA copies of LASVpv was determined to be 1 × 10^7^ copies/ml by using a standard curve generated with plasmids carrying the VSVΔG-Rluc. The HTS assay conditions, including the seeding cell density and LASVpv infective dose, were optimized at 1 × 10^4^ cells and 1 × 10^2^ copies per 96-well plate, respectively. Under the optimized conditions, signal-to-basal (S/B) ratio, coefficient of variation (CV), and Z’ factor were 41770, 11.9%, and 0.62, respectively, demonstrating that the assay was promising for large-scale screening of inhibitors.

The HTS schematic is depicted in Fig. 1A. Inhibitors were defined as primer candidates with inhibition > 50% and no apparent cytotoxicity in duplicate wells at a concentration of 10 μM. Of the 1018 tested compounds, 52 (5.11%) were considered primer candidates. A screening to reconfirm the results was then carried out using these primer candidates over a broader concentration range (3.125 to 50.0 μM). Seven compounds (0.69%) were selected based on their concentration-dependent inhibitory effects and a cell viability > 80%. Subsequently, these 7 compounds were subjected to counter-screening to rule out inhibitors of VSV genome replication and Rluc. Using these criteria, 2 hits, lacidipine and phenothrin, were selected with specific inhibition against LASV GPC activity, while the other 5 compounds were eliminated (Fig. 1B).

**Fig 1.**
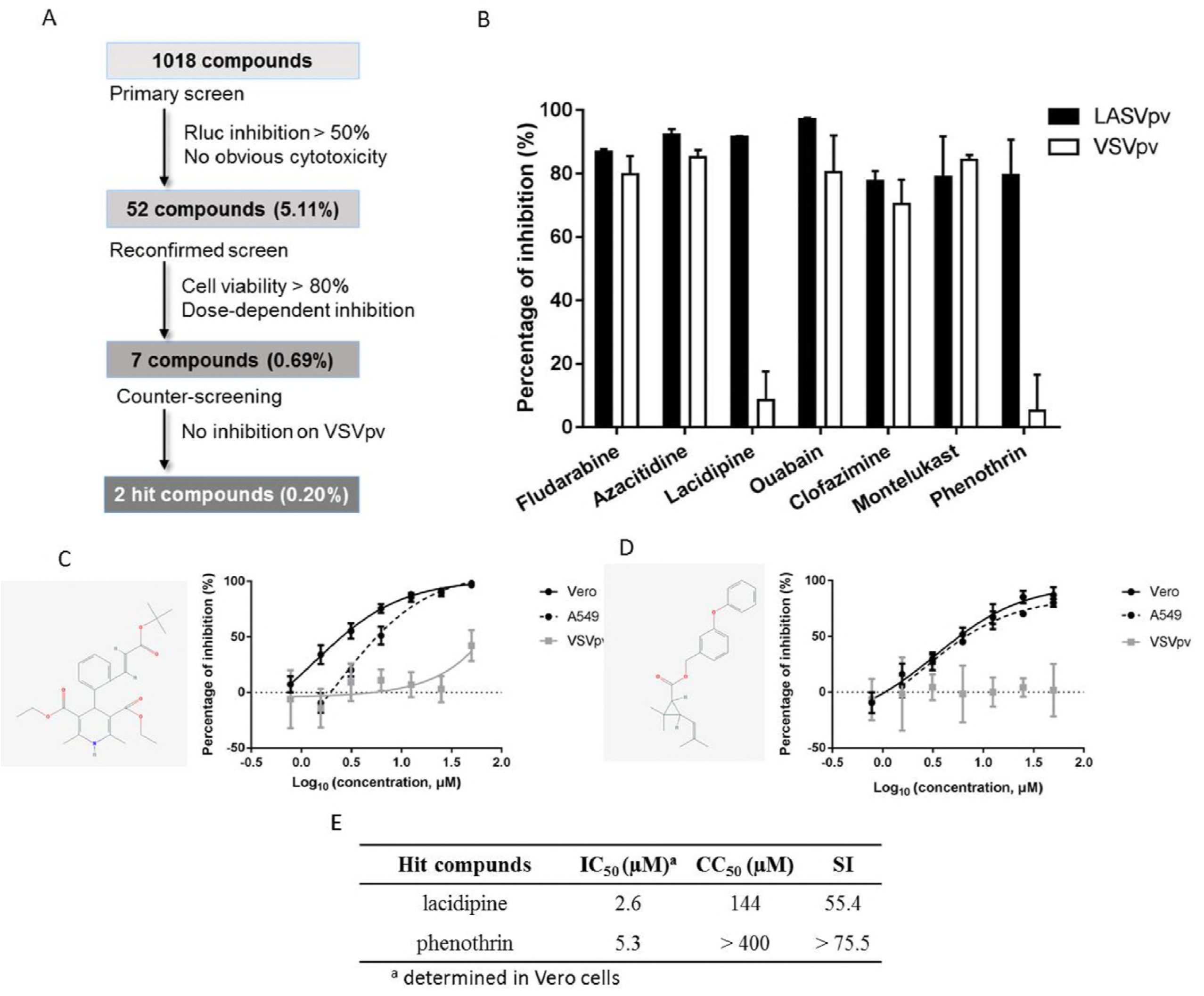
High throughput screening (HTS) for inhibitors of Lassa virus (LASV) entry from a Food and Drug Administration (FDA)-approved drug library. (A) The HTS assay flowchart is shown. (B) A counter-screening of the 7 selected compounds to reconfirm the initial screening results is shown. Vero cells were seeded at a density of 1×10^4^ cells per well in 96-well plates. After incubating overnight, cells were treated in duplicate with compounds (25 μM); pseudo-type LASV (LASVpv) was added 1 h later, with a multiplicity of infection (MOI) of 0.01. The supernatant was removed after 1 h, and the cells were re-treated with the compounds for an additional 23 h. (C and D) Dose-response curves of lacidipine (C) and phenothrin (D) for inhibiting LASVpv infection are shown; the insets in each graph shown the drug structures. (E) The 50% inhibitory concentration (IC_50_), 50% cytotoxic concentration (CC_50_), and selective index (SI) for lacidipine and phenothrin are shown. *Rluc*, Renilla luciferase; VSVrv, recombinant vesicular stomatitis virus; VSVg, pseudo-type vesicular stomatitis virus; IC_50_, 50% inhibitory concentration; CC_50_, 50% cytotoxic concentration; SI, selective index

Lacidipine is a dihydropyridine voltage-gated Ca^2+^ channel antagonist, while phenothrin is a synthetic pyrethroid used for aerosol insecticides. We evaluated the 50% inhibitory concentration (IC_50_) and 50% cytotoxic concentration (CC_50_) of both hit drugs. Both lacidipine and phenothrin exhibited dose-dependent inhibition of LASVpv infections. Additionally, both drugs inhibited LASVpv infection in the A549 human epithelial cell line; epithelial cells are important targets of infection *in vivo*, suggesting these drugs are potentially useful in the treatment of human infections (Fig. 1C and D). The selective index (SI, the ratio of the CC_50_ to the IC_50_) for lacidipine was 55.4, while that for phenothrin was > 75.5 (Fig. 1E). The CC_50_ values for the 2 hit drugs were similar to those previously published for diverse cell systems; however, they were determined using different toxicity assays (25). To validate the antiviral effects, lacidipine and phenothrin were purchased from other commercial sources and tested; the cytotoxic and antiviral effects were similar to the results of our primary screening.

### Lacidipine and Phenothrin Inhibited GPC-mediated Membrane Fusion

Arenavirus GPCs have a unique structure in which the cleaved SSP is retained and non-covalently associates with GP2; many arenavirus entry inhibitors have been shown to bind with and stabilize the prefusion forms of GPC to prevent membrane fusion (26-28); therefore, we asked whether the 2 hit drugs act via a similar mechanism. To address this, 293T cells transfected with GPC were incubated with either drug and subjected to a low-pH pulse to promote fusion. As shown in Fig. 2A, a low pH in the GPC-transfected cells induced obvious membrane fusion, whereas a neutral pH had no effect. Both drugs inhibited syncytium formation at all tested concentrations, suggesting that both drugs inhibit GPC conformational changes induced by an acidic environment.

**Fig 2.**
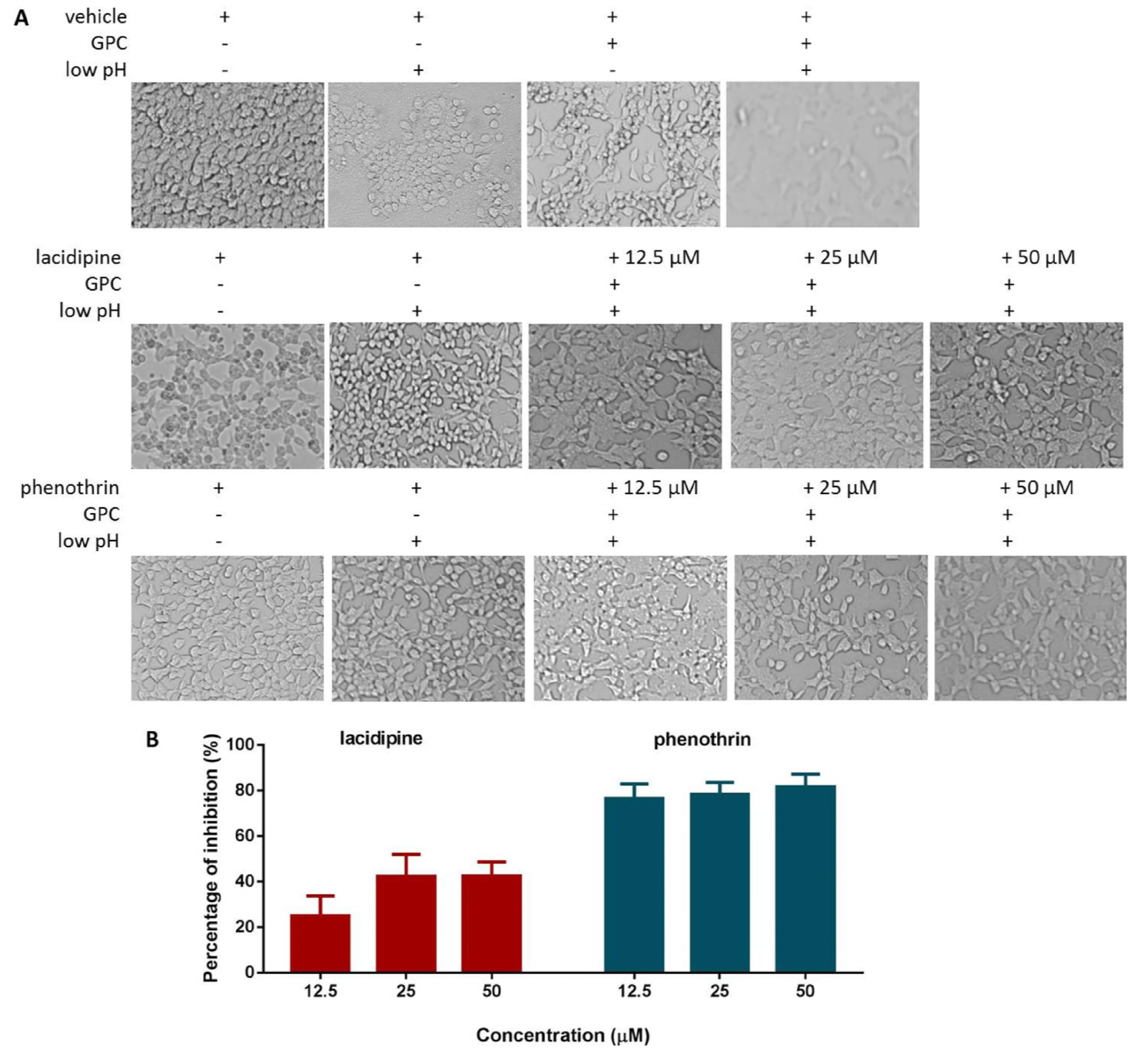
Lacidipine and phenothrin inhibit glycoprotein complex (GPC)-mediated membrane fusion. (A) 293T cells were transfected with pCAGGS-Lassa virus (LASV) GPC or the empty pCAGGS expression plasmid; 24 h later the drugs or vehicle (dimethyl sulfoxide, DMSO) were added for 1 h followed by treatment with acidified (pH 5.0) Dulbecco’s modified Eagle’s medium (DMEM) for 15 min. The cells were then washed and placed in neutral pH DMEM. Syncytium formation was visualized after 3 h using light microcopy. Images are representative fields from 4 to 5 independent experiments. (B) A dual luciferase assay was used to quantitatively evaluate the inhibitory activities of lacidipine and phenothrin against membrane fusion. 293T cells transfected with both pCAGGS-LASV GPC and plasmid expressing pCAGT7 were co-cultured at a ratio of 1:3 with targeted cells transfected with pT7EMCVLuc together with the pRL-CMV control vector. The cell fusion activity was quantitatively determined by measuring firefly luciferase activity and standardized with *Renilla* luciferase (Rluc) activity. Data are presented as means ± SDs for 3 independent experiments.

To further quantitatively evaluate the inhibitory activities, fusion efficacy was determined using a dual-luciferase assay. As shown in Fig. 2B, the maximum inhibitory rates for lacidipine and phenothrin were 42.4% and 81.2%, respectively, at the range of concentrations tested. Notably, phenothrin exhibited great activity (~80%) against GPC-mediated membrane fusion even at the lowest concentration tested (12.5 μM). Together, these results show that both drugs, especially phenothrin, prominently inhibit GPC-mediated membrane fusion.

### Lacidipine Irreversibly Binds to GPC

Lacidipine and phenothrin inhibited GPC-mediated membrane fusion; therefore, we asked whether the drugs irreversibly bind to GPC and prevent conformational changes. To test this, we conducted a virucidal assay to investigate the binding ability of the hit drugs to native GPC. LASVpv was mixed with drugs for 1 h; the mixture was then diluted 200-fold and added to the cells for 1 h. As shown in Fig. 3A, luciferase activity was not suppressed in the phenothrin group, indicating that phenothrin did not irreversibly bind to the neutral-pH forms of GPC (29). However, it is possible that phenothrin binds to a structural GPC intermediate resulting from the low pH. A reduction > 70% was observed in the lacidipine group, suggesting that lacidipine irreversibly interacts with the pre-fusion conformation of GPC and inhibits LASV entry.

**Fig 3.**
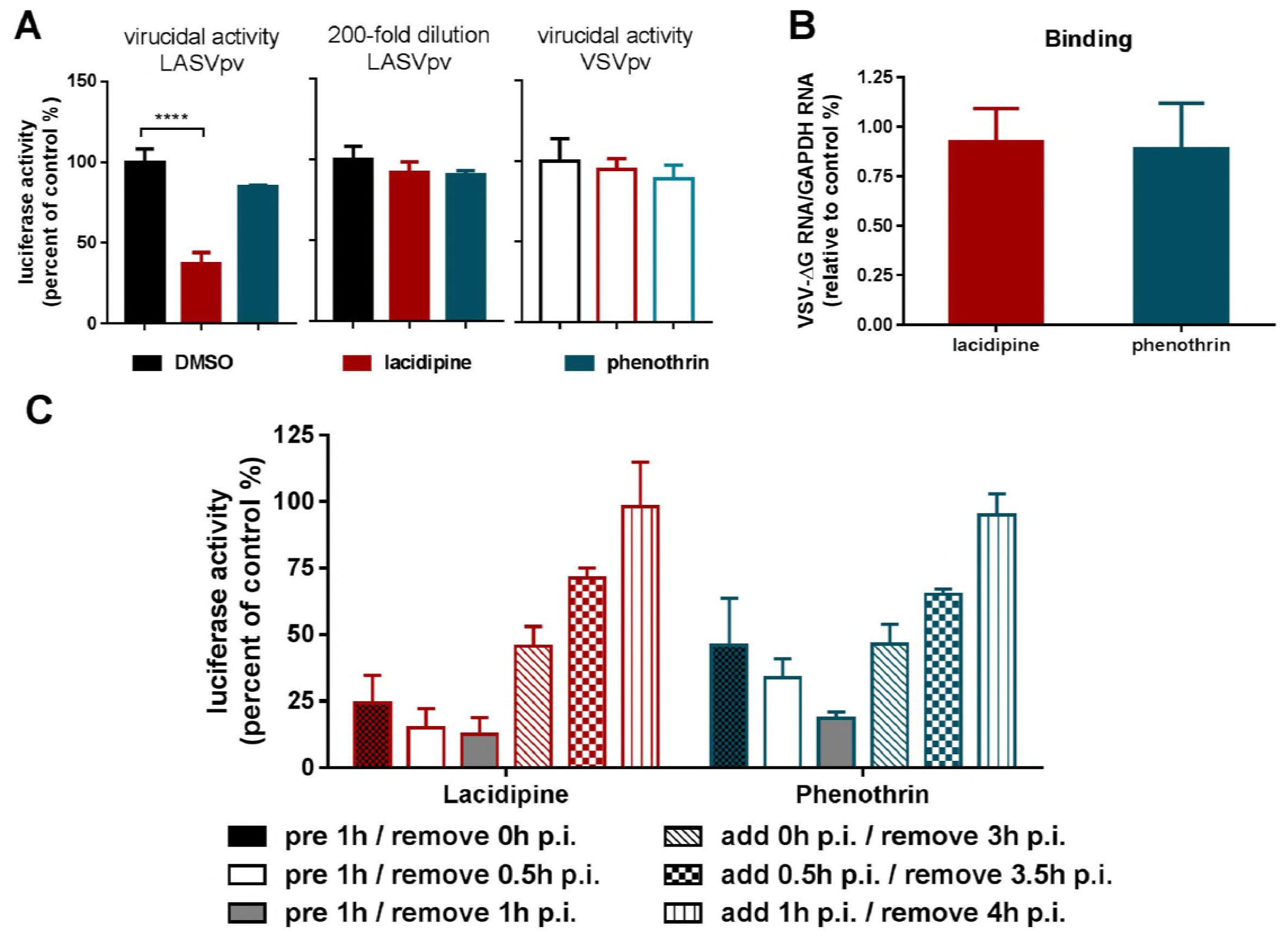
Effects of lacidipine and phenothrin on different stages of Lassa virus (LASV) entry. (A) Results of a virucidal assay are shown. (Left) Pseudo-type LASVs (LASVpv) with a multiplicity of infection (MOI) of 2 were incubated with dimethyl sulfoxide (DMSO) or drugs (25 μM for 1 h) and then diluted 200-fold and added to cells. (Middle) Cells were incubated with DMSO or drugs for 1 h at 0.125 μM before adding LASVpv (MOI of 0.01). (Right) The pseudo-type vesicular stomatitis virus (VSVpv, MOI of 2) was incubated with DMSO or drug (25 μM for 1 h) and then diluted 200-fold before adding to the cells; luciferase activity was determined 24 h later. (B) The effects of drugs on LASVpv binding are shown. Vero cells were incubated with drugs (50 μM) or vehicle at 37 °C for 1 h, followed by incubation with LASVpv (MOI of 10) in the presence or absence of drugs at 4 °C for an additional 1 h. After extensively washing with cold phosphate-buffered saline (PBS), the bound virus was quantified via RT-qPCR. (C) Vero cells were infected with LASVpv (MOI of 0.01) at 4 °C for 1 h and then washed with PBS; the temperature was then increased to 37°C in the presence of lacidipine (10 μM) or phenothrin (25 μM) for the indicated times. Data are presented as the means ± SDs for 4 independent experiments. GAPDH, glyceraldehyde 3-phosphate dehydrogenase, ** *P* < 0.01.

We next investigated the inhibitory effects of the drugs on binding, which is initialized when GP1, the receptor binding subunit, recognizes the primary receptor α-DG (18). Binding efficacy was evaluated in the absence and presence of the drugs, and no significant decrease in the number of LASVpv particles bound to the cell surface at 4°C was observed in either drug-treated group (Fig. 3B), suggesting that neither drug interferes with the receptor binding subunit GP1 (38,39).

To further confirm drug mechanisms, we studied the inhibition kinetics of lacidipine and phenothrin. Pretreatment of cells with lacidipine or phenothrin sharply decreased LASVpv infection even when removed at 30 min post-infection. The addition of drugs 30 min post-infection resulted in a mild inhibitory effect, while the addition of drugs 1 h post-infection had no effect, suggesting that membrane fusion occurred within 1 h; these results agree with previously published results (30,31) confirming that the early stage of infection is the sensitive drug phase.

### SSP T40K Mutation Confers Resistance to Lacidipine

To identify the viral target of the drugs, we selected an adaptive mutant virus by serially passaging LASVrv in the presence of 10 μM lacidipine and 25 μM phenothrin, respectively. Parallel passaging of LASVrv in dimethyl sulfoxide (DMSO) was used as a control. In the lacidipine-treated group, robust resistance was detected after 12 rounds of passaging. When viruses were treated with 10 μM lacidipine, the viral titer after 12 rounds of passaging was about 100-fold higher than that in the wild type (WT) virus (Fig. 4A). The lacidipine-resistant virus isolate was plaque purified and sequenced for the entire glycoprotein precursor GPC region. An amino acid substitution was observed in the isolated clone, but markedly absent in the DMSO-treated virus, resulting in a threonine (T) to lysine (K) switch at amino acid position 40 in the ectodomain of SSP (i.e. the last position of the SSP ectodomain) (Fig. 4B). Notably, in the phenothrin-treated group, no obvious improvement in resistance was detected after 20 rounds of passaging, suggesting that phenothrin was less prone to induce adaptive mutations in the glycoprotein.

**Fig 4.**
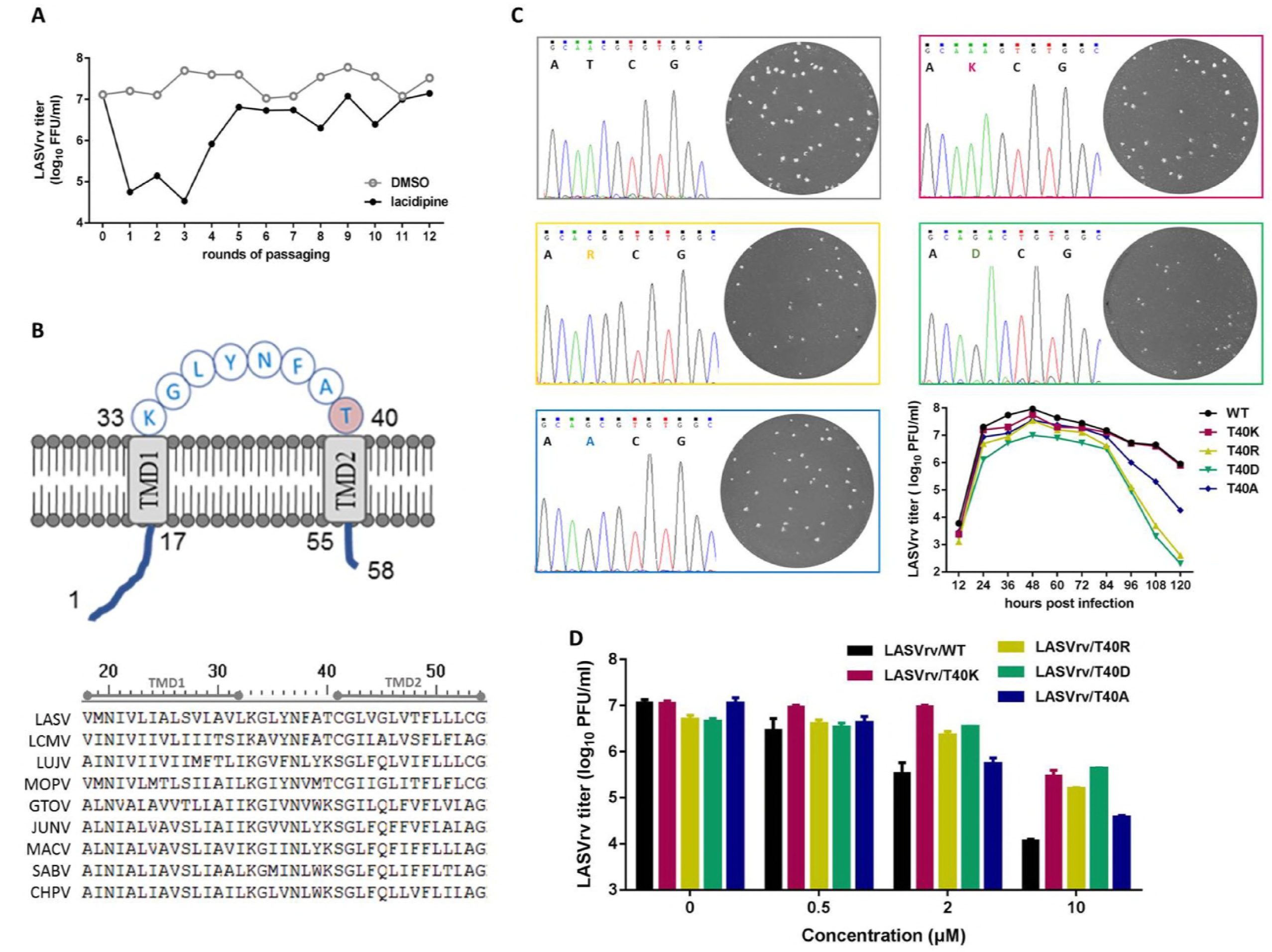
Selection and characterization of lacidipine-resistant recombinant Lassa virus (LASVrv). (A) The adaptive mutant virus was selected by serially passaging LASVrv in the presence of 10 μM lacidipine. In a parallel experiment, LASVrv passaging in vehicle served as a control. After 12 rounds of passaging, no further improvement in resistance was detected, and the selection was terminated. Virus titers and lacidipine sensitivities were determined via a plaque assay in Vero cells. (B) (Top) Membrane topology of LASV stable-signal peptide (SSP) with the threonine 40 (T40) location highlighted is shown (8). (Bottom) The amino acid sequence alignment of the mammarenavirus SSP is shown. The GenBank accession numbers are listed in Materials and Methods. (C) (Top) A sequencing chromatogram of the wild type (WT) and recombinant viruses with plaque morphology of each virus as an inset is shown. (Right bottom) Growth kinetics of the recombinant viruses with different T40 mutations are shown. Vero cells were infected with a multiplicity of infection (MOI) of 0.1 for 1 h. The supernatants were collected at indicated time points post-infection and assayed for the viral titer. Data are presented as means ± SD from 2 independent wells. (D) Resistant activity of the recombinant viruses to lacidipine is shown. Data are presented as means ± SD from 2 independent experiments. GTOV, Guanarito virus; JUNV, Junín virus, MACV, Machupo virus, SABV Sabiá virus; CHPV, Chandipura virus; LCMV, lymphocytic choriomeningitis virus; LUJV, Lujo virus; MOPV, Mopeia virus; LASV, Lassa virus; PFU, plaque forming units; FFU, focus-forming units

The arenavirus SSP, unlike other enveloped viruses, is unusually long and retains the GPC as a vital subunit, playing an essential role in glycoprotein maturation and viral infectivity (12,16,32). The 58-amino acid SSP contains two hydrophobic domains linked by an 8-amino acid ectodomain loop that interacts with the proximal- and trans-membrane region of GP2 to confer sensitivity of fusion inhibitors (26,33,34). To confirm that the T40K mutation conferred lacidipine resistance and to investigate the role of T40 in SSP function and lacidipine inhibition of LASV entry, we produced recombinant viruses with T40K, T40R, T40D, or T40A mutations by introducing the desired mutations into the GPC gene and generating mutant viruses. To investigate the biological properties of the mutant viruses, we first examined the growth kinetics of the rescued viruses. As shown in Fig. 4C, all the viruses caused an accumulation of infectious virions that reached the highest titer at 48 h p.i. Infection of T40K mutant viruses resulted in similar growth curves to those of the WT virus, while T40R and T40D mutants produced less virus after 18 to 36 h. Plaque morphology analyses revealed that the T40K plaques were similar to the WT plaques, whereas T40R and T40D plaques were smaller, and T40A plaques were mid-range. These results suggested that the infectivity of SSP T40R and T40D mutant viruses is milder than that of the WT virus, and position 40 in SSP was tolerable for K.

We next investigated sensitivity of the four mutant viruses to lacidipine. As shown in Fig. 4D, the T40K, T40R, and T40D mutant viruses conferred resistance to lacidipine, which efficiently inhibited LASVrv WT infection at a concentration of 10 μM and reduced viral yields by 3 log units. In contrast, T40K, T40R, and T40D mutant viruses were resistant to lacidipine, with the viral titer decreasing slightly less than 1 log unit; the T40A mutant virus showed no resistance to lacidipine.

Taken together, these results suggest that the T40 mutant was not only critical in conferring lacidipine sensitivity, but also important for LASV infectivity. Substitution of T with K conferred resistance to lacidipine without apparent loss of growth, while substitution with a small nonpolar amino acid (A, alanine) did not affect lacidipine sensitivity. Other positively charged amino acids (R, arginine) or negatively charged amino acids (D, aspartic acid) resulted in the mutant viruses replicating more slowly than the WT virus.

### Lacidipine Affects Entry of Other Arenaviruses

T40 is conserved in OW viruses, except for Lujo virus (LUJV), whereas K40 is similarly conserved (K or R) in NW viruses; therefore, we investigated the effects of lacidipine on the entry of other pathogenic arenaviruses, such as OW viruses (including lymphocytic choriomeningitis virus (LCMV), LUJV, and the closely related Mopeia virus (MOPV) and NW viruses (including JUNV, MACV, GTOV, SABV, and Chandipura virus (CHPV) using a pseudo-type virus. Vero cells were treated with vehicle (DMSO) or lacidipine (1.5625 to 25 μM) starting 1 h before infection (multiplicity of infection [MOI] = 0.01) to 1 h post-infection. At 23 h p.i., cell lysates were collected, and luciferase activity determined. As shown in Fig. 5A, all viruses mentioned above remained unaffected, except for GTOV and MOPV, which exhibited dose-dependent inhibition with an IC_50_ of 6.2 and 4.8 μM, respectively. We also investigated the broad-spectrum antiviral activity of phenothrin, which dose-dependently inhibited the entry of GTOV, MOPV, and CHPV, with IC_50_ values of 6.1, 8.3, and 8.0 μM, respectively. Phenothrin had a less powerful effect on the entry of JUNV, MACV, and SABV, since the percentage of inhibition was less than 50% at the highest tested concentration.

**Fig 5.**
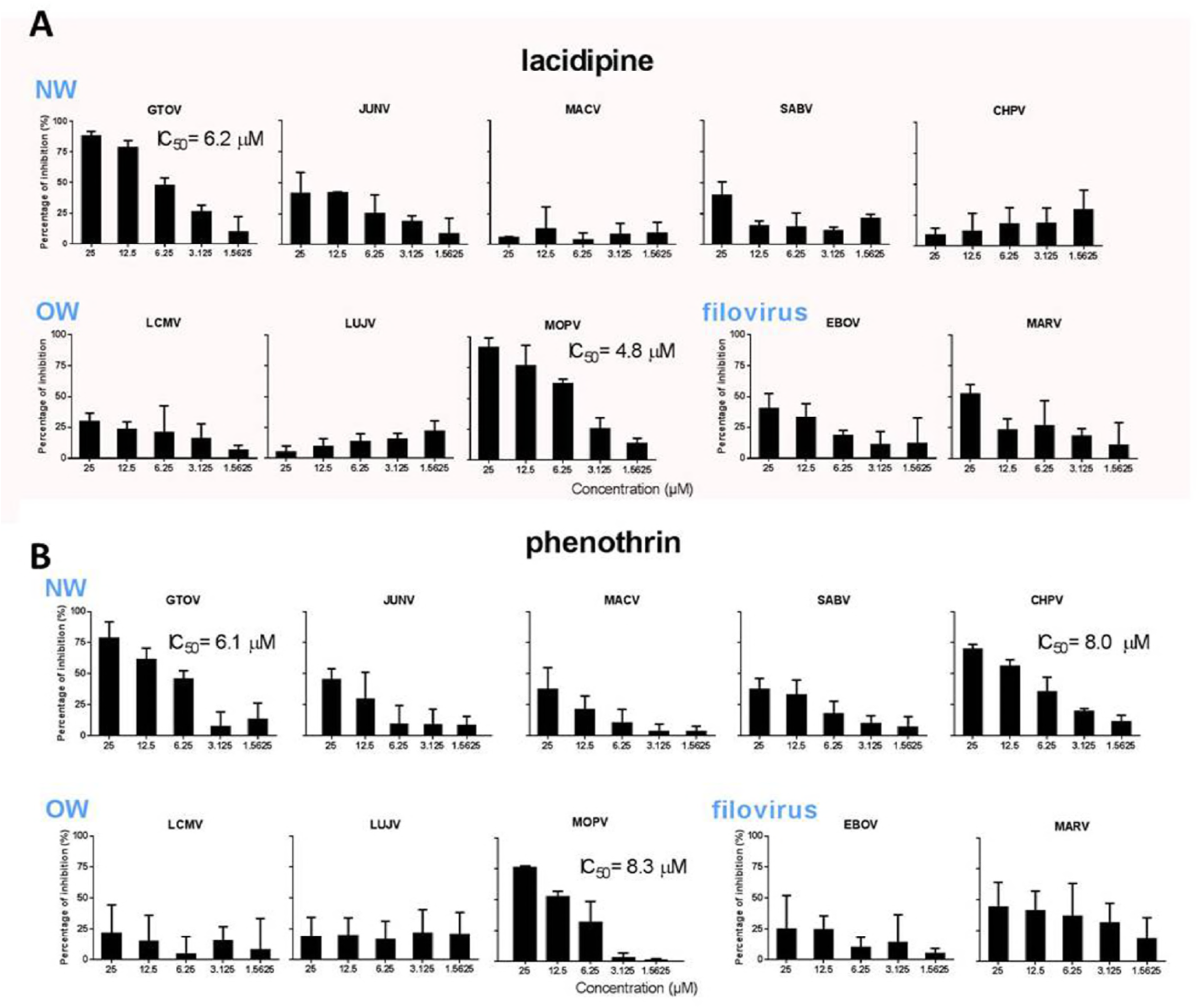
Broad spectrum antiviral activity of the hit drugs against different mammarenavirus and filovirus. Vero cells were incubated in the absence and presence of lacidipine (A) or phenothrin (B). After 1 h, a pseudo-type of the Guanarito virus (GTOV), Junín virus (JUNV), Machupo virus (MACV), Sabiá virus (SABV), Chandipura virus (CHPV), lymphocytic choriomeningitis virus (LCMV), Lujo virus, Mopeia virus (MOPV), Ebola virus (EBOV), and Marburg virus (MARV) were added. The supernatant was removed 1 h later and the cell lysates were assessed for luciferase activity after 23 h. Data are presented as means ± SD from 5 independent experiments. NW, new world; OW, old world

To further assess the role of lacidipine and phenothrin on other class I fusion proteins, we utilized pseudo-type Ebola virus (EBOV) and Marburg virus (MARV). It is important to note that neither lacidipine nor phenothrin treatment robustly inhibited the entry of EBOV and MARV (Fig. 6), indicating a lacidipine- or phenothrin-related interaction with GPC of LASV and GTOV.

### Selection of Lacidipine-resistant GTOVrv

GTOV possesses a K at position 40 of the SSP; therefore, we suggest that an amino acid other than K40 contributed to the sensitivity of GTOV to lacidipine. We further determined the viral target by selecting the adaptive mutant virus by serially passaging GTOVrv in the presence of 10 μM lacidipine. After 15 rounds, 2 amino acid substitutions, V36M in the ectodomain of SSP and V436A in the transmembrane domain of GP2, were observed in the resistant virus (Fig. 6A). To investigate the functional significance of these residues, a pseudo-type GTOV containing the mutants was used to evaluate lacidipine sensitivity.

**Fig 6.**
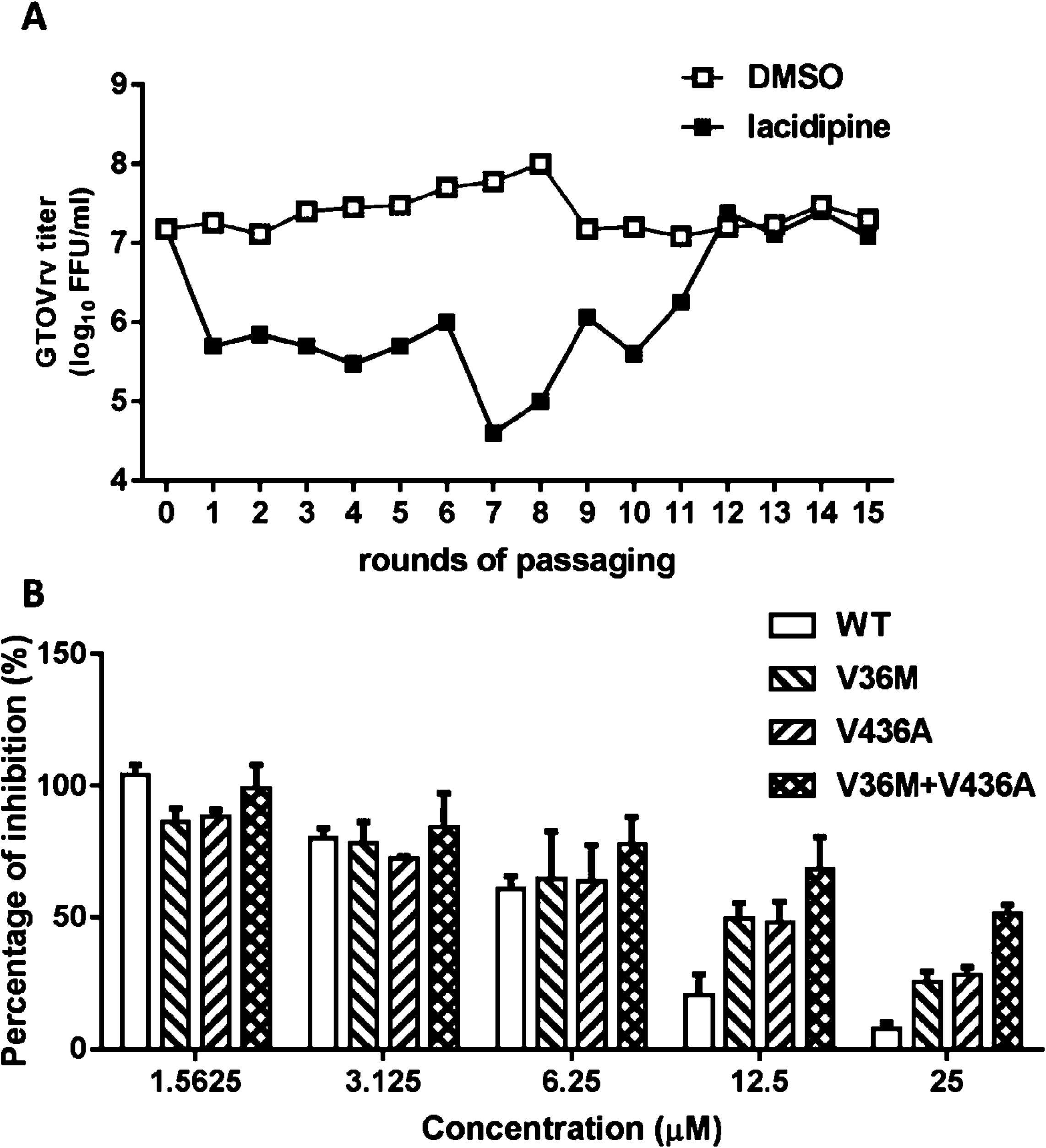
Fig 6 Selection of lacidipine-resistant recombinant Guanarito virus (GTOVrv). (A) The adaptive mutant virus was selected by serially passaging GTOVrv in the presence of 10 μM lacidipine. GTOVrv passaging in vehicle served as a control in parallel. After 15 rounds of passaging, no further improvement in resistance was detected and the selection was terminated. Virus titers and lacidipine sensitivities were determined via plaque assay in Vero cells. (B) Resistant activity of the recombinant viruses to lacidipine is shown. Data are presented as means ± SD from 2 independent experiments. DMSO, dimethyl sulfoxide; WT, wild type; FFU, focus forming units.

GTOV was much less sensitive when either the V36M or V436A mutant was generated, and lacidipine sensitivity was further reduced when both sites were changed (Fig. 6B). As V36 is similarly conserved in NW pathogenic viruses, we reasoned that residues other than these selective mutations contributed to the sensitivity of LASV and GTOV to lacidipine.

## DISCUSSION

In this report, we screened an FDA-approved drug library and identified 2 hit drugs, lacidipine and phenothrin, which prohibited the entry step of LASV infection.

Lacidipine is a lipophilic dihydropyridine calcium antagonist. Since calcium channels proved to be a therapeutic target for other enveloped viruses and calcium inhibitors showed promising effects on the entry of the closely-related JUNV and EBOV (22,42-44), we investigated whether lacidipine inhibits LASV entry by acting as a calcium inhibitor. To address this, we first reviewed all 22 calcium inhibitors included in the current FDA drug library; results showed that in addition to lacidipine, only two calcium inhibitors moderately inhibited LASV entry, including diltiazem and cilnidipine. Both drugs exerted less than 70% inhibition on LASVpv infection at the highest concentration tested (20 μM), suggesting that calcium inhibitors do not effectively block LASV entry as observed for other enveloped viruses. Moreover, lacidipine irreversibly binds to LASV GPC and prohibits acid-pH-induced conformational changes of GPC and subsequent membrane fusion, supporting the speculation that under these conditions, lacidipine functions via a novel mechanism rather than as a calcium inhibitor. To further elucidate the underlying mechanisms associated with prevention of LASV entry via lacidipine, we characterized the viral target of lacidipine by serially passaging LASVrv in the presence of lacidipine. In particular, we observed an amino acid substitution that resulted in a T-to-K switch at amino acid position 40 in the ectodomain of SSP. The SSP in the arenavirus GPC differs from that of conventional signal peptides as it: (i) is unusually long, containing 58 amino acids; (ii) is retained after cleavage, and non-covalently associates with GP2 and GP1 to constitute the GPC heterotrimer; and (iii) interacts with GP2, including the ecto-, transmembrane, and intracellular domains. As a result, the native structure of GPC is stabilized, participates in GPC-mediated activity, and provides an interface targeted by some entry inhibitors (8,12,16,32-40). Among the 8 amino acid residues in the ectodomain of SSP, the absolutely conserved K33 and N37 are well studied and have proven to be essential in GPC maturation, fusion, and infectivity (12,41). In the current study, we demonstrate that replacement of T40 with a charged amino acid (K, R, or D) confers LASV resistance to lacidipine, and position 40 is tolerable for K without detectable changes in LASVrv growth kinetics. These results suggest that T40 together with lacidipine might interplay with the residues located in the membrane-proximal ectodomain of GP2, thus stabilizing the native structure of GPC. Although position 40 is tolerable for K, the stabilization interaction between T40 and lacidipine is collapsed by the replacement of K.

As T40 and K40 are relatively conserved in OW and NW viruses, respectively, we further investigated the broad-spectrum inhibition of lacidipine against other pathogenic mammarenaviruses. As anticipated, MOPV, the most phylogenetic-related virus, was sensitive to lacidipine with a similar IC_50_ value. Intriguingly, LCMV, the OW prototype possessing T at position 40, showed resistance to the drug. Moreover, GTOV, a NW mammarenavirus, was sensitive to lacidipine. Mapping the viral target of GTOV revealed two mutant sites, one in the ectodomain of SSP and the other in the transmembrane domain of GP2. These results suggest the SSP and proximal-membrane ectodomain of GP2 for arenavirus exhibit relatively high amino acid sequence conservation, and the overall GP1-GP2 structure of LASV aligns well with LCMV (42,43); however, the interplay between SSP and GP2 involves a multiple sequential correlation and is not limited at the point-to-point sub interface. Lacidipine-sensitivity depends on accessibility of the interface to the drug as well as the stability of the drug-GPC complex.

We also identified phenothrin via HTS; phenothrin is a pyrethroid usually used in pesticide products and is effective at inhibiting LASV entry. We demonstrated that phenothrin has activity against MOPV, GTOV, and CHPV, with IC_50_ values lower than 10 μM. The structure of phenothrin is similar to that of the LASV specific entry inhibitor ST-161, which uses a cyclopropyl *N*-acylhydrazone as a scaffold (26,44). In our study, resistant viruses for LASV or GTOV were not successfully generated in the presence of phenothrin, suggesting this drug is less prone to induce adaptive mutations in the glycoprotein. In line with this, phenothrin exhibited little effect against LASVpv infection in the virucidal assay, indicating that phenothrin binds less tightly to GPC than lacidipine. Alternatively, phenothrin could bind to a low-pH-induced intermediate conformation of GPC. By understanding the mechanisms of viral entry, inhibitors could be used to uncover novel drug targets, providing further insight into the pathogenesis of LASV.

In conclusion, the findings reported here provide novel insights into the molecular mechanisms underlying LASV entry and offer new and promising therapeutic possibilities for combating arenavirus infections.

## MATERIALS AND METHODS

### Cells and Viruses

BHK-21, HEK 293T, Vero, HeLa, and A549 cells were cultured in Dulbecco’s modified Eagle’s medium (HyClone, Logan, UT, USA) supplemented with 10% fetal bovine serum (Gibco, Grand Island, NY, USA). The pseudo-type VSV bearing the GPC of LASV (Josiah strain, Genbank HQ688673.1), LCMV (Armstrong strain, GenBank AY847350.1), LUJV (GenBank NC_012776.1), MOPV (GenBank AY772170.1), GTOV (GenBank NC_005077.1), JUNV (XJ13 strain, GenBank NC_005081.1), MACV (Carvallo strain, GenBank NC_005078.1), SABV (GenBank U41071.1), CHPV (GenBank NC_010562.1), EBOV (Mayinga strain, GenBank: EU224440.2), and MARV (YP_001531156.1) were generated as previously reported using the infectious clone for the VSV, Indiana serotype (kindly provided by Yoshiharu Matsuura, Osaka University, Osaka, Japan) (45-47). The recombinant VSV expressing the GPC of LASV and GTOV, in which the appropriate open reading frames for the GPC were cloned into the pVSVΔG-eGFP vectors (Plasmid #31842, addgene), were generated as described previously (48,49). The pseudo-type and recombinant viruses enveloped by LASV GPC are designated LASVpv and LASVrv, respectively.

### Optimization of HTS Assay Conditions

Cell density and MOI were optimized for the HTS assay. Vero cells at different densities (2,500–12,500 cells per well) were infected with a MOI from 0.001 to 1 (10-10^4^ copies per well). The appropriate cell density as well as the dose for LASVpv were selected by comparing the signal-to-basal ratio, the coefficients of variation, and Z’ values under different conditions as previously described (29,50). Methyl-beta-cyclodextrin and DMSO were used as a positive and negative control, respectively.

### HTS Assay of an FDA-Approved Compound Library

A library of 1018 FDA-approved drugs was purchased from Selleck Chemicals (Houston, TX, USA). Compounds were stored as 10 mM stock solutions in DMSO at −80 °C until use. The first round HTS was carried out as shown in Fig. 1A. Briefly, Vero cells were seeded at a density of 1×10^4^ cells per well in 96-well plates. After incubating overnight, cells were treated in duplicate with the compounds (10 μM); 1 h later, cells were infected with LASVpv (MOI of 0.01), and the supernatant was removed 1 h post-infection.

The infected cells were lysed 23 h later, and luciferase activity was measured using the Rluc assay system (Promega, Madison, WI). Primary candidates were identified using criteria of no apparent cytotoxicity and an average >50% inhibition in duplicate wells and then subsequently rescreened via serial dilution in triplicate plates to evaluate the IC_50_ (GraphPad Prism 6). Dose-dependent inhibition and cell viability >80% were the criteria used to select 7 compounds. The 7 compounds were then counter-screened using VSVpv (MOI of 0.01) to rule out VSV genome replication inhibitors and Rluc activity. Compounds specifically blocking LASV entry were considered hit drugs and were evaluated for the CC_50_ and SI.

### Membrane Fusion Assay

293T cells transfected with pCAGGS expression plasmids for LASV GPC or the empty pCAGGS were treated with compounds or vehicle (DMSO) for 1 h, followed by incubation for 15 min with acidified (pH 5.0) medium. The cells were then washed and placed in neutral medium, and syncytium formation was visualized 1 h later via light microcopy.

For quantification of the luciferase-based fusion assay, 293T cells transfected with both pCAGGS-LASV GPC and plasmids expressing T7 RNA polymerase (pCAGT7) were co-cultured at a ratio of 3:1 with targeted cells transfected with pT7EMCVLuc and pRL-CMV (plasmids used in this assay were kindly provided by Yoshiharu Matsuura, Osaka University, Osaka, Japan). Drug treatment and pH induction were conducted as described above. Cell fusion activity was quantitatively determined after 24 h by measuring firefly luciferase activity and was standardized with Rluc activity as previously described (27,51).

### Virucidal Assay

To study the virucidal effects of the drugs, approximately 5 × 10^5^ copies of LASVpv or VSVpv were incubated with drugs (25 μM) or vehicle at 37 °C for 1 h; the mixture was diluted 200-fold to the non-inhibitory concentration (MOI of 0.01) to infect Vero cells. Luciferase activity was determined 24 h later as described above.

### Binding Assay

Vero cells were pretreated with 50 μM lacidipine or phenothrin; after 1 h the cells were transferred onto ice, and LASVpv (MOI of 10) was added for 1 h. After washed with cold PBS for 3 times, the bound viral particles were quantified via RT-qPCR using a specific primer pair to detect the VSVΔG-Rluc (primers 5’- GTAACGGACGAATGTCTCATAA -3’ and 5’- TTTGACTCTCGCCTGATTGTAC -3’). All RNA amplifications were normalized to glyceraldehyde 3-phosphate dehydrogenase (GAPDH) RNA (obtained via PCR with the following primers: 5’- TCCTTGGAGGCCATGTGGGCCAT -3’ and 5’- TGATGACATCAAGAAGGTGGTGAAG -3’).

### Time-of-Addition Assay

We performed a time-of-addition experiment to elucidate which stage of LASV entry was inhibited by the drugs. At time 0, Vero cells were infected with LASVpv (MOI of 0.01) at 4 °C for 1 h and washed with phosphate-buffered saline; the temperature was then increased to 37°C (designated time 0 post infection [p.i.]) to synchronize the infections. Test compounds were incubated with the cells as shown in Fig. 3C.

### Selection of Adaptive Mutants

Drug-resistant viruses were generated by passaging LASVrv on Vero cells in the presence of 10 μM lacidipine or 25 μM phenothrin. LASVrv was passaged in the presence of 2% DMSO in parallel as a control. Passaging in the presence of lacidipine was terminated when no further improvement in resistance was detected. RNA from the resistant viruses was extracted, amplified, and purified for sequencing of the GPC segment. Mutant sites were introduced to recover LASVrv as previously described (52). Virus titers and lacidipine sensitivities were determined by means of a plaque assay in Vero cells.

## ACKNOWLEDGEMENTS

We thank the The Center for Instrumental Analysis and Metrology and the Core Facility and Technical Support, Wuhan Institute of Virology for providing technical assistance. This work was supported by the National Natural Sciences Foundation of China (31670165), the Open Research Fund Program of CAS Key Laboratory of Special Pathogens and Biosafety, Wuhan Institute of Virology, and the Open Research Fund Program of Wuhan National Bio-Safety Level 4 Lab of CAS (NBL2017008).

## REFERENCES

1. Oldstone MB. 2002. Arenaviruses. I. The epidemiology molecular and cell biology of arenaviruses. Introduction. Current topics in microbiology and immunology 262:V−XII.

2. Nunberg JH, York J. 2012. The curious case of arenavirus entry, and its inhibition. Viruses 4:83−101.

3. Buchmeier MJ, de la Torre JC, Peters CJ. 2007. Fields Virology, 4th ed. Lippincott-Raven, Philadelphia.

4. Vela E. 2012. Animal models, prophylaxis, and therapeutics for arenavirus infections. Viruses 4:1802−1829.

5. Houlihan C, Behrens R. 2017. Lassa fever. Bmj 358:j2986.

6. Maxmen A. 2018. Deadly Lassa-fever outbreak tests Nigeria’s revamped health agency. Nature 555:421−422.

7. Eichler R, Lenz O, Strecker T, Garten W. 2003. Signal peptide of Lassa virus glycoprotein GP-C exhibits an unusual length. FEBS letters 538:203−206.

8. Wang W, Zhou Z, Zhang L, Wang S, Xiao G. 2016. Structure-function relationship of the mammarenavirus envelope glycoprotein. Virologica Sinica 31:380−394.

9. Igonet S, Vaney MC, Vonrhein C, Bricogne G, Stura EA, Hengartner H, Eschli B, Rey FA. 2011. X-ray structure of the arenavirus glycoprotein GP2 in its postfusion hairpin conformation. Proceedings of the National Academy of Sciences of the United States of America 108:19967−19972.

10. Lenz O, ter Meulen J, Klenk HD, Seidah NG, Garten W. 2001. The Lassa virus glycoprotein precursor GP-C is proteolytically processed by subtilase SKI-1/S1P. Proceedings of the National Academy of Sciences of the United States of America 98:12701−12705.

11. York J, Nunberg JH. 2009. Intersubunit interactions modulate pH-induced activation of membrane fusion by the Junin virus envelope glycoprotein GPC. Journal of virology 83:4121−4126.

12. Saunders AA, Ting JP, Meisner J, Neuman BW, Perez M, de la Torre JC, Buchmeier MJ. 2007. Mapping the landscape of the lymphocytic choriomeningitis virus stable signal peptide reveals novel functional domains. Journal of virology 81:5649−5657.

13. Eichler R, Lenz O, Strecker T, Eickmann M, Klenk HD, Garten W. 2004. Lassa virus glycoprotein signal peptide displays a novel topology with an extended endoplasmic reticulum luminal region. The Journal of biological chemistry 279:12293−12299.

14. Eichler R, Lenz O, Strecker T, Eickmann M, Klenk HD, Garten W. 2003. Identification of Lassa virus glycoprotein signal peptide as a trans-acting maturation factor. EMBO reports 4:1084−1088.

15. Froeschke M, Basler M, Groettrup M, Dobberstein B. 2003. Long-lived signal peptide of lymphocytic choriomeningitis virus glycoprotein pGP-C. The Journal of biological chemistry 278:41914−41920.

16. Agnihothram SS, York J, Trahey M, Nunberg JH. 2007. Bitopic membrane topology of the stable signal peptide in the tripartite Junin virus GP-C envelope glycoprotein complex. Journal of virology 81:4331−4337.

17. Schrempf S, Froeschke M, Giroglou T, von Laer D, Dobberstein B. 2007. Signal peptide requirements for lymphocytic choriomeningitis virus glycoprotein C maturation and virus infectivity. Journal of virology 81:12515−12524.

18. Cao W, Henry MD, Borrow P, Yamada H, Elder JH, Ravkov EV, Nichol ST, Compans RW, Campbell KP, Oldstone MB. 1998. Identification of alpha-dystroglycan as a receptor for lymphocytic choriomeningitis virus and Lassa fever virus. Science 282:2079−2081.

19. Kunz S, Rojek JM, Perez M, Spiropoulou CF, Oldstone MB. 2005. Characterization of the interaction of lassa fever virus with its cellular receptor alpha-dystroglycan. Journal of virology 79:5979−5987.

20. Jae LT, Raaben M, Herbert AS, Kuehne AI, Wirchnianski AS, Soh TK, Stubbs SH, Janssen H, Damme M, Saftig P, Whelan SP, Dye JM, Brummelkamp TR. 2014. Lassa virus entry requires a trigger-induced receptor switch. Science 344:1506−1510.

21. McCormick JB, King IJ, Webb PA, Scribner CL, Craven RB, Johnson KM, Elliott LH, Belmont-Williams R. 1986. Lassa fever. Effective therapy with ribavirin. The New England journal of medicine 314:20−26.

22. Wang S, Liu Y, Guo J, Wang P, Zhang L, Xiao G, Wang W. 2017. Screening of FDA-Approved Drugs for Inhibitors of Japanese Encephalitis Virus Infection. Journal of virology 91:e01055−01017.

23. Chong CR, Sullivan DJ, Jr. 2007. New uses for old drugs. Nature 448:645−646.

24. Garbutt M, Liebscher R, Wahl-Jensen V, Jones S, Moller P, Wagner R, Volchkov V, Klenk HD, Feldmann H, Stroher U. 2004. Properties of replication-competent vesicular stomatitis virus vectors expressing glycoproteins of filoviruses and arenaviruses. Journal of virology 78:5458−5465.

25. Nagy K, Racz G, Matsumoto T, Adany R, Adam B. 2014. Evaluation of the genotoxicity of the pyrethroid insecticide phenothrin. Mutation research. Genetic toxicology and environmental mutagenesis 770:1−5.

26. York J, Dai D, Amberg SM, Nunberg JH. 2008. pH-induced activation of arenavirus membrane fusion is antagonized by small-molecule inhibitors. Journal of virology 82:10932−10939.

27. Thomas CJ, Casquilho-Gray HE, York J, DeCamp DL, Dai D, Petrilli EB, Boger DL, Slayden RA, Amberg SM, Sprang SR, Nunberg JH. 2011. A specific interaction of small molecule entry inhibitors with the envelope glycoprotein complex of the Junin hemorrhagic fever arenavirus. The Journal of biological chemistry 286:6192−6200.

28. Shankar S, Whitby LR, Casquilho-Gray HE, York J, Boger DL, Nunberg JH. 2016. Small-molecule fusion inhibitors bind the pH-sensing SSP-GP2 subunit interface of the Lassa virus envelope glycoprotein. Journal of virology.

29. Rathbun JY, Droniou ME, Damoiseaux R, Haworth KG, Henley JE, Exline CM, Choe H, Cannon PM. 2015. Novel Arenavirus Entry Inhibitors Discovered by Using a Minigenome Rescue System for High-Throughput Drug Screening. Journal of virology 89:8428−8443.

30. Moraz ML, Pythoud C, Turk R, Rothenberger S, Pasquato A, Campbell KP, Kunz S. 2013. Cell entry of Lassa virus induces tyrosine phosphorylation of dystroglycan. Cellular microbiology 15:689−700.

31. Oppliger J, Torriani G, Herrador A, Kunz S. 2016. Lassa virus cell entry via dystroglycan involves an unusual pathway of macropinocytosis. Journal of virology.

32. Bederka LH, Bonhomme CJ, Ling EL, Buchmeier MJ. 2014. Arenavirus stable signal peptide is the keystone subunit for glycoprotein complex organization. mBio 5:e02063.

33. Larson RA, Dai D, Hosack VT, Tan Y, Bolken TC, Hruby DE, Amberg SM. 2008. Identification of a broad-spectrum arenavirus entry inhibitor. Journal of virology 82:10768−10775.

34. Shankar S, Whitby LR, Casquilho-Gray HE, York J, Boger DL, Nunberg JH. 2016. Small-Molecule Fusion Inhibitors Bind the pH-Sensing Stable Signal Peptide-GP2 Subunit Interface of the Lassa Virus Envelope Glycoprotein. Journal of virology 90:6799−6807.

35. Albarino CG, Bird BH, Chakrabarti AK, Dodd KA, White DM, Bergeron E, Shrivastava-Ranjan P, Nichol ST. 2011. Reverse genetics generation of chimeric infectious Junin/Lassa virus is dependent on interaction of homologous glycoprotein stable signal peptide and G2 cytoplasmic domains. Journal of virology 85:112−122.

36. Messina EL, York J, Nunberg JH. 2012. Dissection of the role of the stable signal peptide of the arenavirus envelope glycoprotein in membrane fusion. Journal of virology 86:6138−6145.

37. Burri DJ, Pasquato A, da Palma JR, Igonet S, Oldstone MB, Kunz S. 2013. The role of proteolytic processing and the stable signal peptide in expression of the Old World arenavirus envelope glycoprotein ectodomain. Virology 436:127−133.

38. Shao J, Liu X, Ly H, Liang Y. 2016. Characterization of the glycoprotein stable signal peptide in mediating Pichinde viral replication and virulence. Journal of virology.

39. Briknarova K, Thomas CJ, York J, Nunberg JH. 2011. Structure of a zinc-binding domain in the Junin virus envelope glycoprotein. The Journal of biological chemistry 286:1528−1536.

40. Lee AM, Rojek JM, Spiropoulou CF, Gundersen AT, Jin W, Shaginian A, York J, Nunberg JH, Boger DL, Oldstone MB, Kunz S. 2008. Unique small molecule entry inhibitors of hemorrhagic fever arenaviruses. The Journal of biological chemistry 283:18734−18742.

41. York J, Nunberg JH. 2006. Role of the stable signal peptide of Junin arenavirus envelope glycoprotein in pH-dependent membrane fusion. Journal of virology 80:7775−7780.

42. Hastie KM, Zandonatti MA, Kleinfelter LM, Heinrich ML, Rowland MM, Chandran K, Branco LM, Robinson JE, Garry RF, Saphire EO. 2017. Structural basis for antibody-mediated neutralization of Lassa virus. Science 356:923−928.

43. Hastie KM, Igonet S, Sullivan BM, Legrand P, Zandonatti MA, Robinson JE, Garry RF, Rey FA, Oldstone MB, Saphire EO. 2016. Crystal structure of the prefusion surface glycoprotein of the prototypic arenavirus LCMV. Nature structural & molecular biology.

44. Burgeson JR, Gharaibeh DN, Moore AL, Larson RA, Amberg SM, Bolken TC, Hruby DE, Dai D. 2013. Lead optimization of an acylhydrazone scaffold possessing antiviral activity against Lassa virus. Bioorganic & medicinal chemistry letters 23:5840−5843.

45. Tani H, Shiokawa M, Kaname Y, Kambara H, Mori Y, Abe T, Moriishi K, Matsuura Y. 2010. Involvement of ceramide in the propagation of Japanese encephalitis virus. J Virol 84:2798−2807.

46. Zhang LK, Xin QL, Zhu SL, Wan WW, Wang W, Xiao G. 2016. Activation of the RLR/MAVS signaling pathway by the L protein of Mopeia virus. Journal of virology.

47. Whitt MA. 2010. Generation of VSV pseudotypes using recombinant DeltaG-VSV for studies on virus entry, identification of entry inhibitors, and immune responses to vaccines. Journal of virological methods 169:365−374.

48. Geisbert TW, Jones S, Fritz EA, Shurtleff AC, Geisbert JB, Liebscher R, Grolla A, Stroher U, Fernando L, Daddario KM, Guttieri MC, Mothe BR, Larsen T, Hensley LE, Jahrling PB, Feldmann H. 2005. Development of a new vaccine for the prevention of Lassa fever. PLoS medicine 2:e183.

49. Safronetz D, Mire C, Rosenke K, Feldmann F, Haddock E, Geisbert T, Feldmann H. 2015. A recombinant vesicular stomatitis virus-based Lassa fever vaccine protects guinea pigs and macaques against challenge with geographically and genetically distinct Lassa viruses. PLoS neglected tropical diseases 9:e0003736.

50. Anantpadma M, Kouznetsova J, Wang H, Huang R, Kolokoltsov A, Guha R, Lindstrom AR, Shtanko O, Simeonov A, Maloney DJ, Maury W, LaCount DJ, Jadhav A, Davey RA. 2016. Large-Scale Screening and Identification of Novel Ebola Virus and Marburg Virus Entry Inhibitors. Antimicrobial agents and chemotherapy 60:4471−4481.

51. Takikawa S, Ishii K, Aizaki H, Suzuki T, Asakura H, Matsuura Y, Miyamura T. 2000. Cell fusion activity of hepatitis C virus envelope proteins. Journal of virology 74:5066−5074.

52. Liu H, Liu Y, Wang S, Zhang Y, Zu X, Zhou Z, Zhang B, Xiao G. 2015. Structure-based mutational analysis of several sites in the E protein: implications for understanding the entry mechanism of Japanese encephalitis virus. J Virol 89:5668−5686.

